# Exploring the preservation of a parasitic trace in decapod crustaceans using finite elements analysis

**DOI:** 10.1101/2023.12.07.570666

**Authors:** Nathan Wright, Adiël A. Klompmaker, Elizabeth Petsios

## Abstract

The fossil record of parasitism is poorly understood, due largely to the scarcity of strong fossil evidence of parasites. Understanding the preservation potential for fossil parasitic evidence is critical to contextualizing the fossil record of parasitism. Here, we present the first use of X-ray computed tomography (CT) scanning and finite elements analysis (FEA) to analyze the impact of a parasite-induced fossil trace on host preservation. Four fossil and three modern decapod crustacean specimens with branchial swellings attributed to an epicaridean isopod parasite were CT scanned and examined with FEA to assess differences in the magnitude and distribution of stress between normal and swollen branchial chambers. The results of the FEA show highly localized stress peaks in reaction to point forces, with higher peak stress on the swollen branchial chamber for nearly all specimens and different forces applied, suggesting a possible shape-related decrease in the preservation potential of these parasitic swellings. Broader application of these methods as well as advances in the application of 3D data analysis in paleontology are critical to understanding the fossil record of parasitism and other poorly represented fossil groups.

## Introduction

Parasitism is the most common and ecologically impactful mode of life today, as parasitic taxa comprise at least 40% of described extant species and are critical to ecosystem trophic structure [1]. However, evidence of parasitic behavior in the fossil record is uncommon [2]. Studies describing cases of potential fossil evidence of parasites span several decades [3-5], but broader synthesis and interpretation of fossil parasitic evidence has only recently been undertaken because of broader data access, an increase in studies, and computationally intensive tools [6]. Despite these advances, existing evidence and data of fossil parasitism is limited, taxonomically biased, and often over-represented by sites of exceptional preservation. Taphonomic bias is suggested to be a main factor causing these disparities because many ubiquitous modern parasitic taxa are soft-bodied, small, and/or endoparasitic, with little chance of producing recognizable fossil evidence [7-8]. Many modern marine parasites create distinct traces on their host’s skeletal elements, yet, for many taxa, fossil evidence of these parasites and traces is uncommon. Biases against evidence of parasitism in the fossil record have largely yet to be critically examined or quantified [9-13]. As a result of the scarcity of parasitic evidence, the fossil record of parasitism remains a major gap in the scientific understanding of the evolutionary and ecological history of life on earth.

Crustaceans are a highly diverse, abundant, and globally distributed group of arthropods, with a fossil record spanning more than 500 million years. Crustaceans are well documented as a group in which many novel parasitic interactions have arisen independently several times, with crustaceans exhibiting great diversity as both hosts and parasites [12]. Some of the parasitic relationships among crustaceans produce distinct, preservable traces, that can be identified from fossil host specimens [12]. These features make crustaceans an excellent model group for studying the fossil record of parasitism. The fossil record of crustaceans is limited, and often fragmentary, owing to a lightly to moderately (and often heterogeneously) calcified exoskeleton that readily disarticulates in the absence of soft tissue [14]. Many crustaceans are found in concretions, typically calcite or siderite, which preserve high fidelity 3D fossils, sometimes with original exoskeletal material remaining [15-17]. Fossiliferous concretions are a unique and exceptional taphonomic scenario, suggesting rapid burial and rapid microbially and geochemically mediated concretionary growth of carbonate minerals, which only occurs under specific conditions [18]. Crustaceans have also been the subject of taphonomic study, revealing the fragmentary and biased nature of the crustacean fossil record, controlled largely by differences in calcification across individuals and between taxa and depositional conditions [9,14,19].

Extant crustacean hosts and their parasites are the subject of much study, as a result of being relatively ubiquitous in marine ecosystems, as well as their significant economic and ecological importance, but fossil evidence of parasitism among crustaceans is limited [12]. Suboval branchial swellings found on decapod carapace fossils, identified as the ichnotaxon *Kanthyloma crusta* [11], are among the most well understood examples of fossil parasitism of crustaceans. These traces are inferred to be induced by parasitic epicaridean isopods that commonly produce identical traces on modern decapods [11]. These distinct fossil swellings have been found on diverse decapod fossils since an early peak in the number of parasitized host species in the Late Jurassic, found predominantly in Europe [20]. Likely using the Tethys as a dispersal pathway, *K. crusta* has been observed on decapod fossils globally through the Cretaceous and Cenozoic, although with lower host diversity than seen in the Late Jurassic [21]. Extant isopods in the family Bopyridae, which induce branchial swellings attributable to *K. crusta*, are globally widespread and infest diverse decapod species [22], in contrast to the relatively poor Cenozoic fossil record of *K. crusta*. Modern bopyrid isopods are widespread parasites of decapods, typically at relatively low prevalence (<5%) within host populations, but can instigate rapid host population collapse when introduced to a naïve host population [13, 23].

As high-quality specimens of fossil decapods are rare and scientifically valuable, particularly those bearing additional abnormalities such as traces of parasitism, destructive sampling techniques and intrusive attempts to reveal fossils from the matrix are inadvisable. CT scanning has seen a dramatic increase in widespread use for paleontological study over the last 20 years, as the technology has become more accessible, and CT data has become easier to analyze and manipulate [24-30].

Computational simulation techniques, such as finite element methods and computational fluid dynamics, have been used to simulate physical forces acting on 3D models to explore functional morphology and ecology in fossils [31-32]. Studies over the last two decades have applied finite elements analysis (FEA) to paleontological and zoological material, including morphological study of fossil arthropods and modern crustaceans [32-33], and found that high-fidelity finite-element models of living and fossil organisms are robust at modelling strain and resolving relationships between form and function [34-36].

The aim of this study is to present the first application of FEA in evaluating the impact of the parasite-induced trace fossil *Kanthyloma crusta* on host preservation by exploring how deformations of carapace shape alter the physical preservation potential of the host. Specimens of modern and fossil decapod crustaceans with branchial swellings attributed to isopod infestation (*K. crusta*) were CT-scanned, and FEA was conducted to observe differences in the magnitude and distribution of stress on healthy and swollen branchial chambers.

## Methods

### Specimens studied

Seven specimens of recent and fossil decapod specimens with swellings attributable to *K. crusta* were studied through institutional loans: three Recent specimens of *Munida valida* [37] from the Gulf of Mexico preserved in 70% ethanol (Fig 1 A-C), three complete fossil specimens of *Macroacaena rosenkrantzi* [38] in siderite concretions from the Maastrichtian of Greenland (Fig 1 D-F), and one isolated fossil carapace of *Panopeus nanus* [39] in lower Miocene limestone from the Duncans Quarry, Trelawny Parish, Jamaica (FLMNH-IP locality XJ015) (Fig 1 G). These specimens were studied through institutional loans from Texas A&M University (TAMU), the Natural History Museum of Denmark (NHMD), and the University of Florida, Florida Museum of Natural History, Invertebrate Paleontology (UF-FLMNH-IP), respectively. The recent specimens of *Munida valida* are corpses, the fossil specimens of *Macroacaena rosenkrantzi* likely represent corpses, and the fossil specimen of *Panopeus nanus* likely represents a molt. Each specimen has one swollen branchial chamber, in either the left (n=3), or right (n=4) chamber.

**Fig 1.**
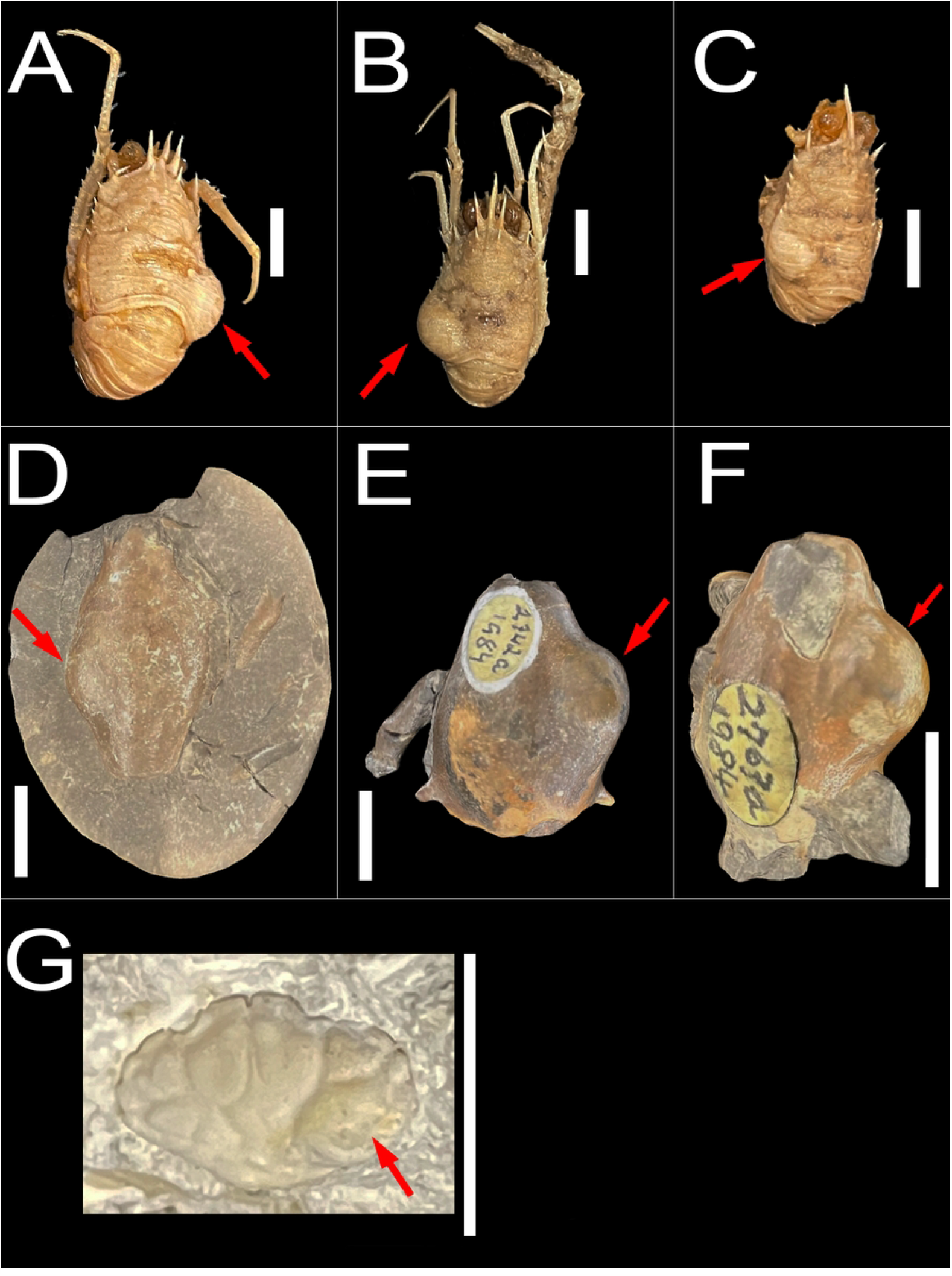
Specimens with branchial swellings. All scale bars are 1cm. Red arrows indicate swellings. A-C) Modern specimens of *Munida valida* TAMU cat. no. 2-3061 (A,C), 2-3063 (B). D-F) Fossil specimens of *Macroacaena rosenkrantzi* NHMD GM 1985.886 (D), GM 1984.2742 (E), GM 1984.2763 (F). G) Fossil specimen of *Panopeus nanus* UF 288470.

### Scanning and CT data preparation

X-ray computed tomography (CT) is a technology that uses an X-ray source to construct several cross-section radiographs, which can then be combined to form a 3D volumetric render of the scanned subject [24]. Specimen 3D data for analysis was captured using a North Star Imaging (NSI) X3000 industrial CT x-ray inspection system. All CT scans were conducted at sub-35μm resolution. Simultaneous CT scans of multiple specimens (including the *Munida* and *Macroacaena* specimens) utilized the NSI SubpiX [40] scanning technique for increased resolution. CT data was initially processed and visualized using the NSI efX software and was then exported as TIFF images stacks for further preparation. The image stacks output was imported into the software Dragonfly ORS 2022.1 [41] for visualization and 3D conversion of the CT data. The specimens were segmented from the raw data using a mixture of manual segmentation and manually trained AI-assisted segmentation models [41]. Segmented specimen data was rendered for visualization in Dragonfly, then exported as Stereolithography (STL) 3D models. Specimen STLs were imported into the software Blender 3.4 [42] for additional cleaning and processing, including removal of unconnected elements, remeshing, downscaling, and fixing geometry errors. After cleaning and fixing errors, each model was standardized to a uniform size and to a uniform number of faces (20,000±1,000) to improve comparability of results between specimens.

### Finite elements analysis

Finite elements analysis (FEA) is a simplified method of testing the load and distribution of physical stresses on a complex 3D model with given material properties, forces, and constraints, by subdividing a complex model into finite parts, then solving partial differential equations for the parts individually [43]. The prepared specimen 3D models were imported into the software FreeCAD 0.20.2 [44] and converted into FEA models using the program Gmsh 4.11.1 [45]. In FreeCAD, identical material properties were applied to each model, which were amalgamated from multiple studies of the material properties of decapod skeletal elements [46-50]. The ventral surfaces of the specimens were set as constraints, and a force of ten newtons of point load force was applied at perpendicular angles to faces at the same location on both the left and right branchial chamber of each specimen (for each specimen, this included one normal branchial chamber and one swollen branchial chamber) (Fig 2). All FEA was also run using applied forces of five newtons and twenty newtons, to assess results sensitivity. Swanson et al. (2013) [51] demonstrated that ten newtons may be sufficient to indent or fracture a crustacean carapace of similar sizes to the specimens included here. Blue crabs, common scavengers and predators of small crustaceans along the east coast of the Americas, have been shown to exert point load forces in excess of 10 newtons at the tip of the dactyl in vivo [52-53]. The finite elements analysis was solved using the solver CalculiX 2.10 [54], then the results were exported into Paraview 5.11.0 [55] as Visualization Toolkit (VTK) files for observation and visualization of FEA results. Von Mises stress, the stress response of a given material relative to the limit at which the material deforms, was chosen as the primary FEA output metric for characterizing the stress, strain, and deformation of the models in response to a force [56]. The vertex data from the left and right side of each mesh were isolated from each other to compare von Mises stress values between the normal and swollen side of each specimen.

**Fig 2.**
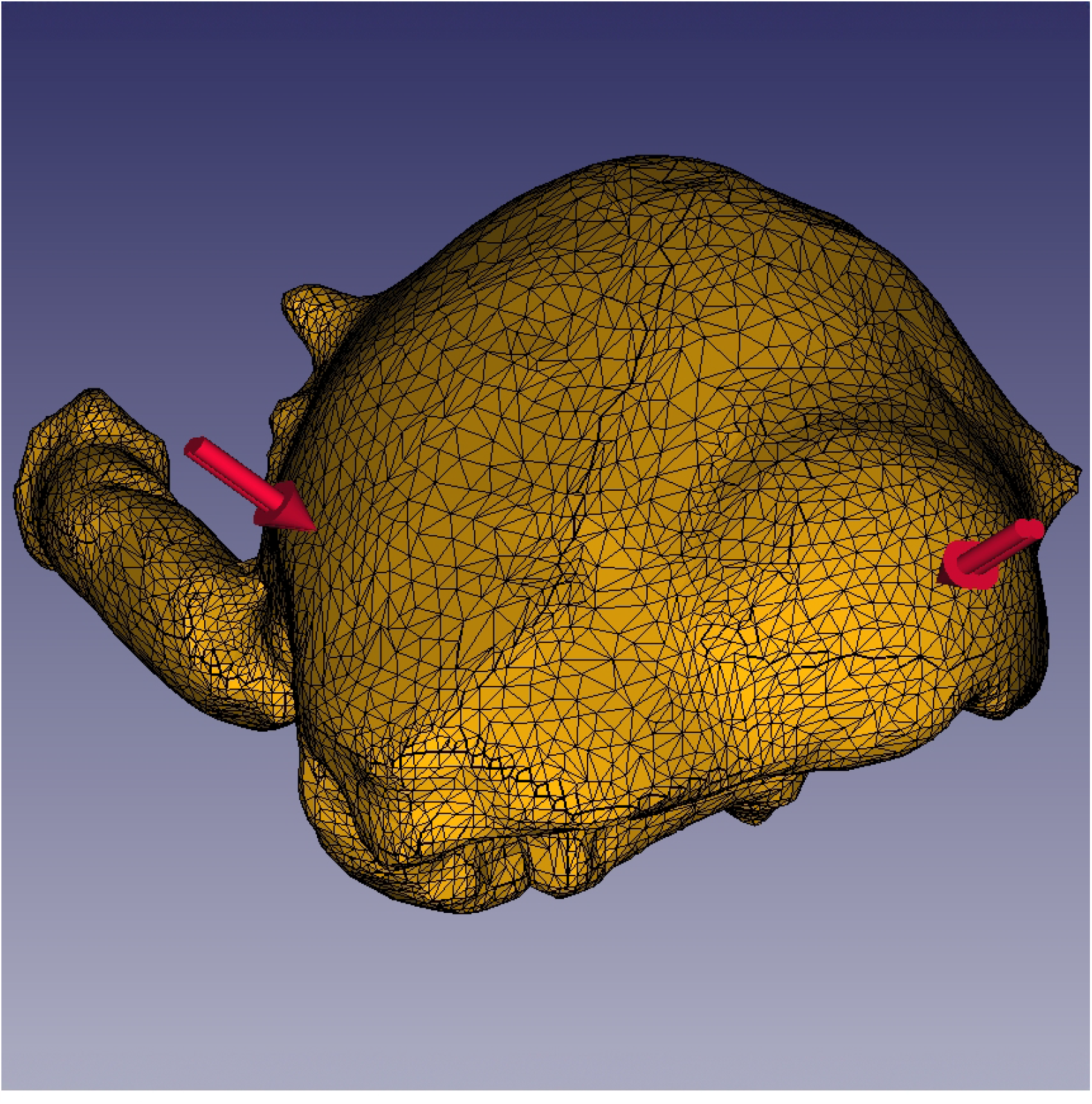
Finite Element Model of *Macroacaena rosenkrantzi* with constraints in FreeCAD. Red arrows indicate applied forces.

## Results

### Computed Tomography

Specimen CT scans output high-resolution 3D data, with a voxel size of 30.5 microns for the specimen of *P. nanus*, and a voxel size of 26.4 microns for the *Munida valida* and *Macroacaena rosenkrantzi* specimens (Fig 3 A-G). The higher resolution of the *Munida* and *Macroacaena* specimens is a result of the SubpiX scanning technique. CT data of the recent *Munida* specimens revealed complex internal morphology and considerable heterogeneity of cuticle density and thickness (Fig 3 A-C). The scans of the fossil *Macroacaena* in concretions had no recognizable internal morphology, or internal evidence of the isopod parasite, although some of the original host cuticle was intact and is distinguishable from the concretion (Fig 3 D-F). The *Panopeus* specimen CT data revealed that it is an isolated carapace, possibly a molt, with no preserved cuticle, and that the swollen right side of the carapace is filled with limestone material, while the normal left side of the carapace contains a large void (Fig 3 G).

**Fig 3.**
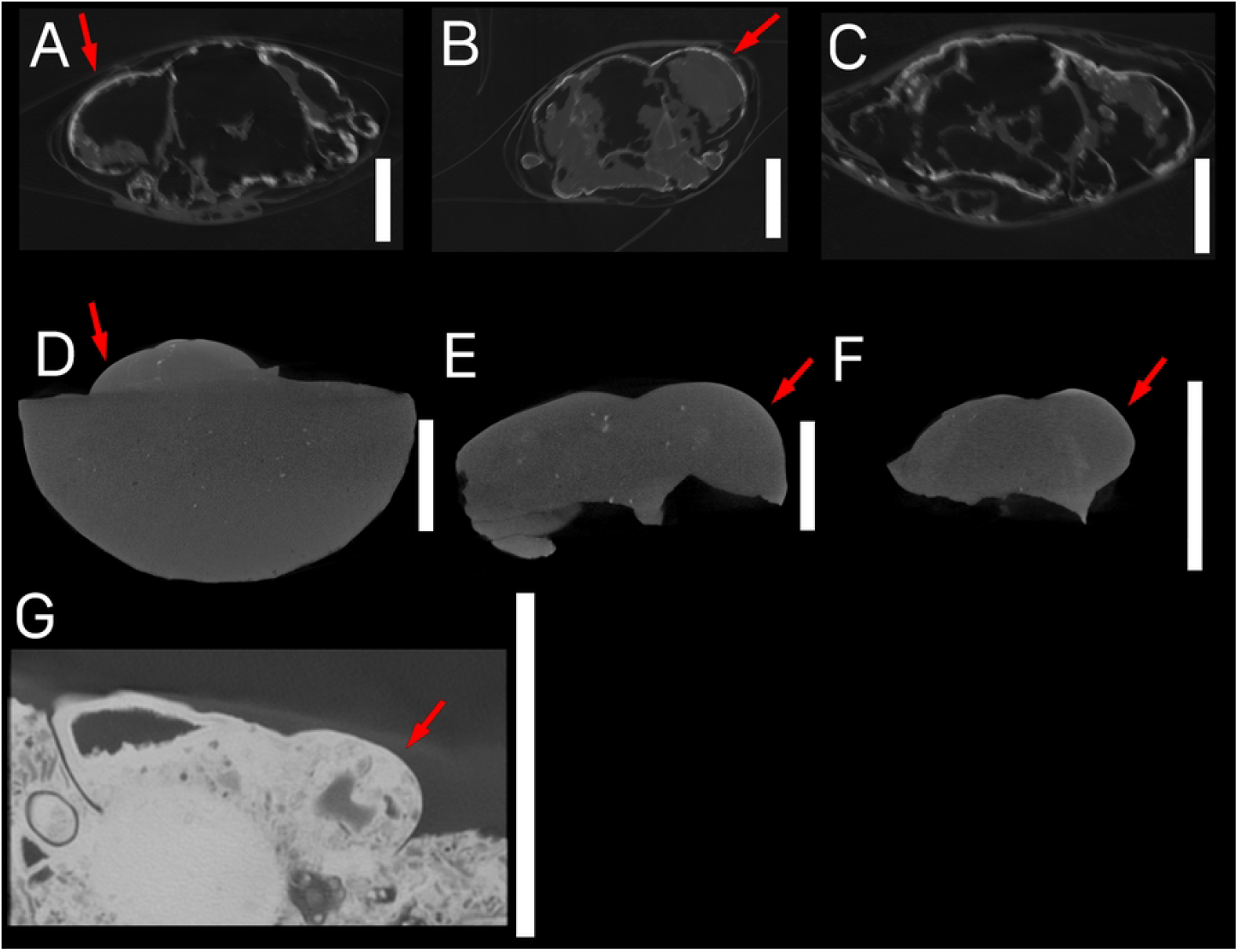
CT data of specimens. All specimens are shown in transverse cross-section. Scale bars are 1cm, shown to the right of each specimen. Red arrows indicate swellings. A-C) Modern specimens of *Munida valida* TAMU cat. no. 2-3061 (A,C), 2-3063 (B). D-F) Fossil specimens of *Macroacaena rosenkrantzi* NHMD GM 1985.886 (D), GM 1984.2742 (E), GM 1984.2763 (F). G) Fossil specimen of *Panopeus nanus* UF 288470.

### Finite Elements Analysis

The finite elements analysis output meshes comprised of tens of thousands of vertices which store individual stress, strain, and deformation values. The stress was highly localized to the area immediately around the site of applied force in all models and was not distributed to other areas of the branchial chamber or carapace (Fig 4). The peak stress on the swollen chamber was higher than on the healthy chamber for all specimens, although the magnitude of the difference varied considerably between specimens, from a 1.53% difference in the specimen of *P. nanus*, to a 51.00% difference in *M. valida* specimen A, with a mean of a 24.13% difference in peak stress, and median of 20.54% peak stress across all specimens (Figs 4 and 5, Table 1). When considering high-stress values directly at the site of the applied force (Fig 5), there are minimal differences in distribution between the normal and swollen sides of each specimen, although the broader trend of highly localized stress peaks, resulting in right-skewed distributions, is observed in most specimens even when excluding smaller stress values. These distributions of stress and differences in peak stress are insensitive to changes in input force, except for the specimen of *P. nanus*, which had peak stress vary between the swollen and healthy chamber at different input forces but maintained peak stress differences below 2% (tested at 5N and 20N, S1 and S2 Figs, S1 Table). FEA results stress values are stored and accessible through the repository Dryad [57].

**Fig 4.**
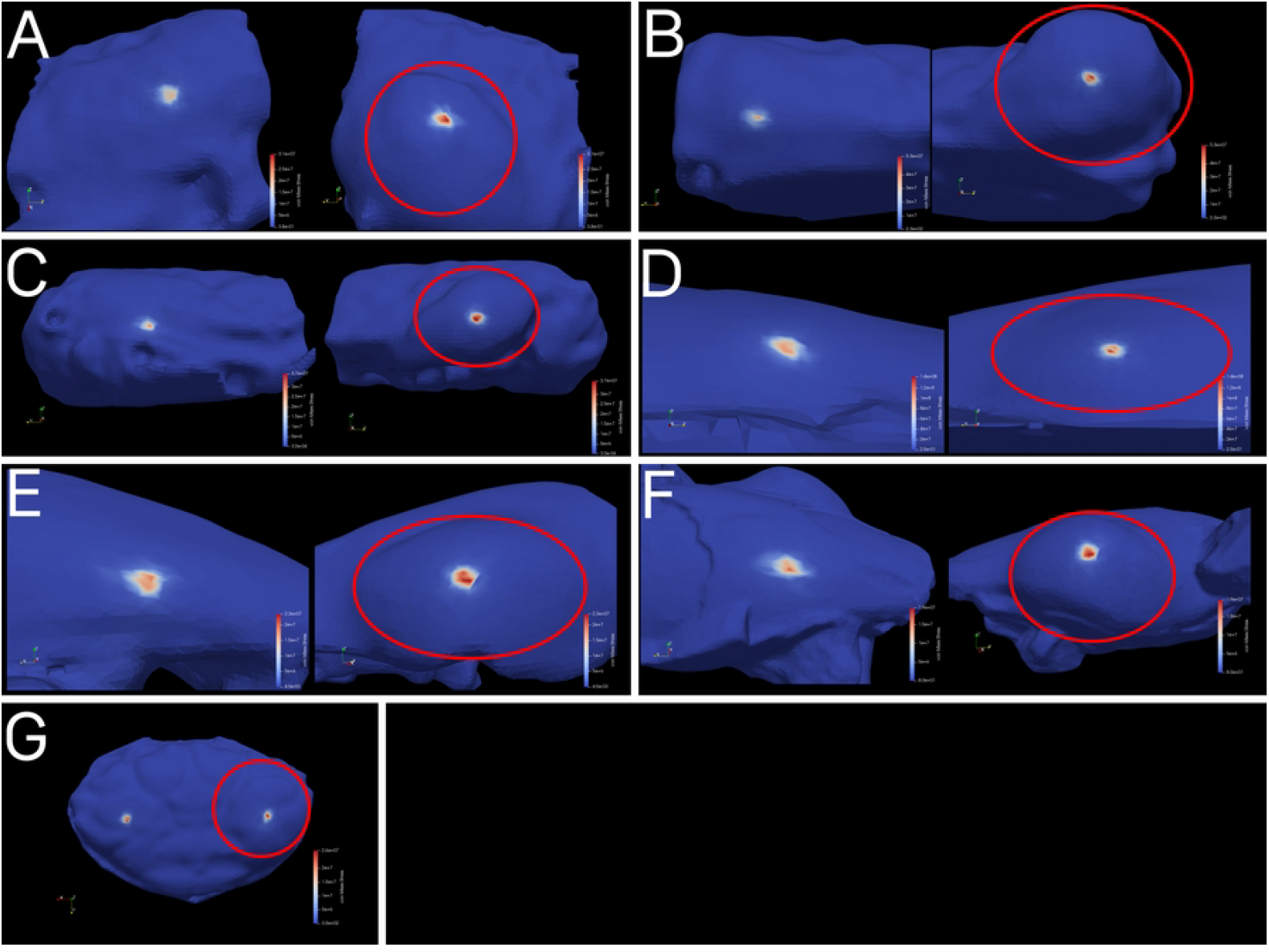
FEA results meshes. Swellings are displayed to the right and circled in red for each specimen. Values on scales are in pascals (Pa). A-C) Modern specimens of *Munida valida* TAMU cat. no. 2-3061 (A,C), 2-3063 (B). D-F) Fossil specimens of *Macroacaena rosenkrantzi* NHMD GM 1985.886 (D), GM 1984.2742 (E), GM 1984.2763 (F). G) Fossil specimen of *Panopeus nanus* UF 288470.

**Fig 5.**
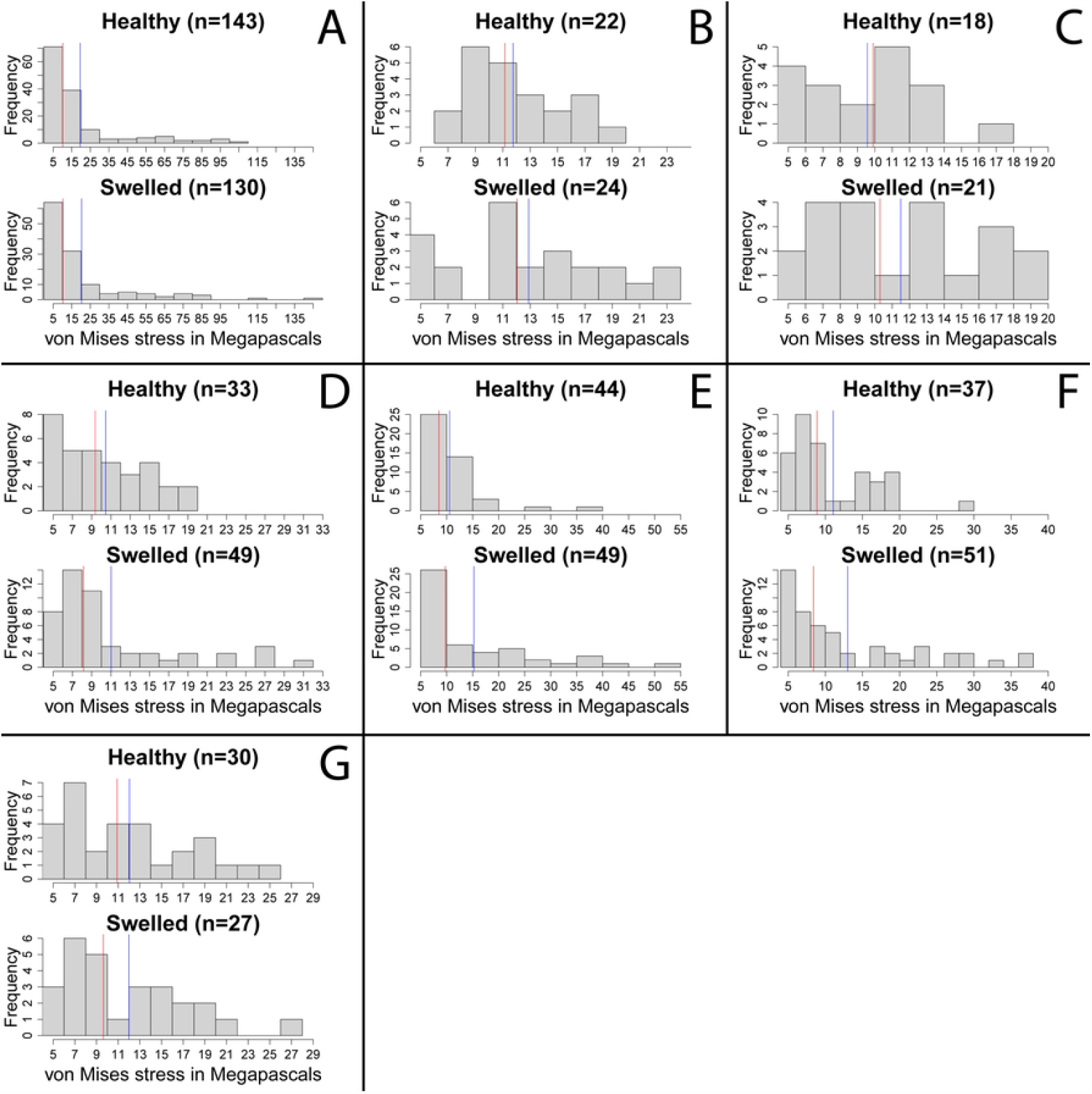
Histograms of model vertex von Mises stress values exceeding 5 MPa. Red lines indicate the median stress values. Blue lines indicate the mean stress values. The number of vertices exceeding 5MPa is given for each side of each specimen. A) *Munida valida* specimen A TAMU cat. no. 2-3061. B) *M. valida* specimen B TAMU cat. No. 2-3063. C) *M. valida* specimen C TAMU cat. no. 2-3061. D) *Macroacaena rosenkrantzi* specimen A NHMD GM 1985.886. E) *M. rosenkrantzi* specimen B NHMD GM 1984.2742. F) *M. rosenkrantzi* specimen C NHMD GM 1984.2763. G) *Panopeus nanus* UF 288470.

**Table 1.** Summary of von Mises stress values from specimen FEA.

|  | Healthy<br>- Peak<br>Stress | Healthy<br>- Mean<br>Stress | Healthy<br>- Mean<br>Stress<br>above<br>5MPa | Healthy -<br>Median<br>Stress<br>above<br>5MPa | Percent<br>Difference<br>in Peak<br>Stress | Swollen<br>- Peak<br>Stress | Swollen<br>- Mean<br>Stress | Swollen -<br>Mean<br>Stress<br>above<br>5MPa | Swollen -<br>Median<br>Stress<br>above<br>5MPa |
| --- | --- | --- | --- | --- | --- | --- | --- | --- | --- |
| *Munida valida* A<br>TAMU cat. no. 2-3061 | 18.7 | 0.0924 | 10.45 | 9.37 | 51.00% | 31.5 | 0.1521 | 11.02 | 8.16 |
| *Munida valida* B<br>TAMU cat. no. 2-3063 | 39.4 | 0.076 | 10.56 | 8.51 | 29.81% | 53.2 | 0.0877 | 15.24 | 9.75 |
| *Munida valida* C<br>TAMU cat. no. 2-3061 | 29.7 | 0.0763 | 11.07 | 8.87 | 20.54% | 36.5 | 0.101 | 13.01 | 8.42 |
| *Macroacaena*<br>*rosenkrantzi* A<br>NHMD GM<br>1985.886 | 102 | 0.1574 | 19.52 | 10.1 | 32.79% | 142 | 0.1425 | 20.36 | 10.2 |
| *Macroacaena*<br>*rosenkrantzi* B<br>NHMD GM<br>1984.2742 | 19.2 | 0.0947 | 11.75 | 11.15 | 20.14% | 23.5 | 0.0953 | 12.89 | 12.05 |
| *Macroacaena*<br>*rosenkrantzi* C<br>NHMD GM<br>1984.2763 | 17.1 | 0.0721 | 9.57 | 9.90 | 13.11% | 19.5 | 0.0675 | 11.49 | 10.3 |
| *Panopeus nanus*<br>UF 288470 | 25.9 | 0.0574 | 12.04 | 10.9 | 1.53% | 26.3 | 0.0516 | 11.99 | 9.64 |

All stress values are in megapascals (MPa).

## Discussion

The results of the finite elements analysis for the seven specimens are indicative of higher overall stress at a swollen branchial chamber relative to a healthy branchial chamber resulting from the same force. Four of the specimens have greater overall mean stress on the swollen chamber relative to the healthy chamber, and all specimens except for the *M. rosenkrantzi* specimen C have greater median stress on the swollen chamber, although due to the highly localized nature of each models’ stress response (Fig 4), and the greater number of vertices on a swollen branchial region resulting from the shape deformity relative to a normal branchial region, means and medians of stress values are not directly comparable. When the results include only the vertices with high stress values around the site of the applied force (above 5 MPa: Fig 5 and Table 1), the number of vertices is similar between healthy and swollen chambers, four of the specimens have greater median stress on the swollen chamber, and the mean stress is higher on the swollen chamber for all but the specimen of *P. nanus*. These patterns, as well as the overall distribution of stress and differences in peak stress, are insensitive to changes in the magnitude of applied force, although the peak stress on the specimen of *P. nanus* varied between the healthy and swollen chamber at different forces while maintaining a peak difference below 2% (S1 and S2 Figs, S1 Table). This difference in *P. nanus*, in combination with the small difference in peak stress between normal and swollen regions and greater variability of results under different forces, is likely a reflection of the considerably different body plan of *Panopeus* relative to *Munida* and *Macroacaena*. The relatively elongated and rounded body plans of *Munida* and *Macroacaena* resulted in laterally exaggerated branchial swellings relative to the flat and wide body plan of *Panopeus*, which has the least visually apparent swelling of the specimens studied. This difference may reflect broader differences in the impact of *K. crusta* on preservation across decapod taxa and body plans.

The observed trend, with greater peak stress in response to the same force at the site of a parasite-induced branchial swelling relative to a normal branchial chamber for each specimen, may represent an early indication of a shape-related decrease in fossil preservation potential for *Kanthyloma crusta*. If correct, a taphonomic bias against the preservation of this parasitic trace implies reduced fossil prevalence and host diversity relative to the fossil record of unparasitized hosts. These results emphasize the importance of recontextualizing the fossil record of *K. crusta* and parasitism more broadly to disentangle fossil record and sampling biases from true signals in the fossil record of parasitism. Non-destructive, computationally intensive methods, as presented here, are key to revealing ecological and evolutionary trends in the fossil record of parasitism.

Resolving this gap is crucial as it has become imperative to understand and predict how a changing climate will impact parasite-host dynamics. Studies predicting changes in parasite-host dynamics relating to anthropogenic change have produced varying results, owing to geographic and taxonomic unevenness of climate change impacts. However, it appears likely that climate change will induce shifts in the range, prevalence, and diversity of parasites, leading to widespread consequences for global ecosystems [58-62]. It is crucial to constrain the fossil record of parasite evolution and ecology to better create a baseline for the impacts of anthropogenic change, as well as to model the consequences of climate change on parasites from comparable climate events in the fossil record, such as the Paleocene-Eocene Thermal Maximum [63].

Fossil preservation is a complex combination of biological, ecological, and environmental factors and conditions which are not fully explored here. Although point load forces as used in this analysis can be applied by some common crustacean scavengers and predators [52-53], point load forces do not represent the full range of forces and pressures that decapod skeletal material may be subjected to before burial. Decapods exhibit considerable differences in shape, cuticle thickness, and calcification between species, as well as ecological differences between taxa which have considerable taphonomic impacts [14]. In addition, the presence of soft tissue and reabsorption of calcium before molting may result in considerable differences in preservation between corpses and molts [64]. Each of these factors must be examined together going forward to understand the diverse preservation characteristics of crustaceans and their parasites through time, and subsequently crustacean parasite-host dynamics throughout earth’s history. In this study of seven specimens across three species, the changes to body shape induced by a crustacean parasite swelling and their impact on preservation were isolated from the many other factors affecting preservation. Thus, there is considerable work still to be done, both in understanding the relationship between shape and preservation in these fossil parasite traces, and in contextualizing the preservation of the fossil record of crustacean parasites more broadly. The methods presented here must be further refined and broadly applied to better characterize the preservation of parasites in the marine fossil record. The complexity of preservation necessitates future refinements to the FEA methods used here, which should strive to incorporate increased model resolution, improved fidelity of material properties, and implementation of carapace thickness, as well as implementing advances to the fidelity and application of FEA in paleontology such as non-linearities [65]. The significant differences between species and individuals also emphasize the importance of investigating patterns of parasite host-preservation with larger numbers of fossil and modern specimens, and across larger numbers of species. These computationally intensive scanning and analytical methods are especially valuable non-destructive tools for in-depth study of rare fossil occurrences, such as parasitic traces, but it is also critical to ground truth these results through experimentation with modern proxies, as well as thorough study of institutional fossil collections.

## Acknowledgments

We thank the following individuals and institutions for their guidance and kind loans of the specimens used in this study: Mary Wicksten and Texas A&M University; Roger Portell and the University of Florida, Florida Museum of Natural History, Invertebrate Paleontology; Laura Cotton, Arden Bashforth, and the Natural History Museum of Denmark. (Reviewers…)

## Supporting Information

**S1 Fig. FEA results meshes for 5N force**. Swellings are displayed to the right and circled in red for each specimen. A-C) Modern specimens of *Munida valida* TAMU cat. no. 2-3061 (A,C), 2-3063 (B). D-F) Fossil specimens of *Macroacaena rosenkrantzi* NHMD GM 1985.886 (D), GM 1984.2742 (E), GM 1984.2763 (F). G) Fossil specimen of *Panopeus nanus* UF 288470.

**S2 Fig. FEA results meshes for 20N force**. Swellings are displayed to the right and circled in red for each specimen. A-C) Modern specimens of *Munida valida* TAMU cat. no. 2-3061 (A,C), 2-3063 (B). D-F) Fossil specimens of *Macroacaena rosenkrantzi* NHMD GM 1985.886 (D), GM 1984.2742 (E), GM 1984.2763 (F). G) Fossil specimen of *Panopeus nanus* UF 288470.

**S1 Table.** Summary of von Mises stress values from specimen FEA at 5N and 20N.

| 5 Newtons | Healthy - Peak Stress | Percent Difference in Peak Stress | Swollen - Peak Stress |
| --- | --- | --- | --- |
| *Munida valida* A TAMU cat. no. 2-3061 | 9.37 | 50.74% | 15.74 |
| *Munida valida* B TAMU cat. no. 2-3063 | 19.71 | 29.72% | 26.59 |
| *Munida valida* C TAMU cat. no. 2-3061 | 14.86 | 20.53% | 18.26 |
| *Macroacaena rosenkrantzi* A NHMD GM 1985.886 | 50.94 | 32.61% | 70.79 |
| *Macroacaena rosenkrantzi* B NHMD GM 1984.2742 | 9.58 | 20.18% | 11.73 |
| *Macroacaena rosenkrantzi* C NHMD GM 1984.2763 | 8.57 | 12.68% | 9.73 |
| *Panopeus nanus* UF 288470 | 22.05 | 1.97% | 21.62 |
| 20 Newtons |  |  |  |
| *Munida valida* A TAMU cat. no. 2-3061 | 37.5 | 50.67% | 62.95 |
| *Munida valida* B TAMU cat. no. 2-3063 | 78.85 | 29.71% | 106.36 |
| *Munida valida* C TAMU cat. no. 2-3061 | 59.44 | 20.52% | 73.03 |
| *Macroacaena rosenkrantzi* A NHMD GM 1985.886 | 203.75 | 32.62% | 283.17 |
| *Macroacaena rosenkrantzi* B NHMD GM 1984.2742 | 38.31 | 20.23% | 46.93 |
| *Macroacaena rosenkrantzi* C NHMD GM 1984.2763 | 34.29 | 12.62% | 38.91 |
| *Panopeus nanus* UF 288470 | 88.18 | 1.96% | 86.47 |

All stress values are in Megapascals (MPa).

## References

1. Dobson A, Lafferty KD, Kuris AM, Hechinger RF, Jets W. Homage to Linnaeus: How many parasites? How many hosts? PNAS. 2008 Aug 12;105(Suppl 1):11482–89.

2. Leung TLF. Fossils of parasites: What can the fossil record tell us about the evolution of parasitism? Biol Rev. 2015 Nov 5;92:410–30.

3. Cressey R, Patterson C. Fossil parasitic copepods from a Lower Cretaceous Fish. Science. 1973 Jun 22;180(4092):1283–5.

4. Baumiller TK. Non-predatory drilling of Mississippian crinoids by platyceratid gastropods. Palaeontology. 1990 Aug;33(3):743–8.

5. Feldmann RM. Parasitic castration of the crab, Tumidocarcinus giganteus Glaessner, from the Miocene of New Zealand: Coevolution within the Crustacea. Journal of Paleontology. 1998 May;72(3):493–8.

6. De Baets K, Huntley JW, Scarponi D, Klompmaker AA, Skawina A. Phanerozoic parasitism and marine metazoan diversity: Dilution versus amplification. Phil. Trans. R. Soc. B. 2021 Sep 20;376:20200366.

7. De Baets K, Littlewood DTJ. Chapter One—The importance of fossils in understanding the evolution of parasites and their vectors. Advances in Parasitology. 2015 Nov 21;90:1–51.

8. Littlewood DTJ, Donovan SK. Fossil parasites: A case of identity. Geology Today. 2003 Jul;19(4):136–42.

9. Klompmaker AA, Portell RW, Frick MG. Comparative experimental taphonomy of eight marine arthropods indicates distinct differences in preservation potential. Paleontology. 2017 Jul 16;60:773–94.

10. Upeniece I. The unique fossil assemblage from the Lode Quarry (Upper Devonian, Latvia). Mitt. Mus. Nat.kd. Bcrl., Geowiss. Reihe. 2001 May 11;4:101–19.

11. Klompmaker AA, Artal P, van Bakel BWM, Fraaije RHB, Jagt JWM. Parasites in the fossil record: A Cretaceous fauna with isopod-infested decapod crustaceans, infestation patterns through time, and a new ichnotaxon. PLoS ONE. 2014 Mar 25;9(3):e92551.

12. Klompmaker AA, Boxshall GA. Chapter Six—Fossil crustaceans as parasites and hosts. Advances in Parasitology. 2015 Jul 10;90:233–89.

13. Dumbauld BR, Chapman JW, Torchin ME, Kuris AM. Is the collapse of mud shrimp (Upogebia pugettensis) populations along the Pacific coast of North America caused by outbreaks of a previously unknown bopyrid isopod parasite (Orthione griffenis)? Estuaries and Coasts. 2010 Jul 9;34:336–50.

14. Bishop GA. Taphonomy of the North American decapods. Journal of Crustacean Biology. 1986 Jul 1;6(3):326–55.

15. McCoy VE, Young RT, Briggs DEG. Factors controlling exceptional preservation in concretions. Palaios. 2015 Apr 1;30(4):272–80.

16. Waugh DA, Feldmann RM, Crawford RS, Jakobsen SL, Thomas KB. Epibiont preservational and observational bias in fossil marine decapods. Journal of Paleontology. 2004 Sep;78(5):961–72.

17. Feldmann RM, Frantescu A, Frantescu OD, Klompmaker AA, Logan G Jr, Robins CM, Schweitzer CE, Waugh DA. Formation of lobster-bearing concretions in the late cretaceous Bearpaw shale, Montana, United States, in a complex geochemical environment. Palaios. 2012 Dec 1;27(12):842–56.

18. Wilson DD, Brett CE. Concretions as sources of exceptional preservation, and decay as a source of concretions: Examples from the Middle Devonian of New York. Palaios. 2013 May 1;28(5):305–16.

19. Krause RA, Parsons-Hubbard K, Walker SE. Experimental taphonomy of a decapod crustacean: Long-term data and their implications. Palaeogeography, Palaeoclimatology, Palaeoecology. 2011 Dec 15;312(3-4):350–62.

20. Klompmaker AA, Robins CM, Portell RW, De Angeli A. Crustaceans as hosts of parasites throughout the phanerozoic. Topics in Geobiology. 2022 Jan 1;50:121–72.

21. Lima D, Alencar DR, Santana W, Oliveira NC, Saraiva AÁF, Oliveira GR, Boyko CB, Pinheiro AP. 110-million-years-old fossil suggests early parasitism in shrimps. Scientific Reports. 2023 Sep 4;13:14549.

22. Williams JD, Boyko CB. The global diversity of parasitic isopods associated with crustacean hosts (Isopoda: Bopyroidea and Cryptoniscoidea). PLoS ONE. 2012 Apr 25;7(4):e35350.

23. Smith AE, Chapman JW, Dumbauld BR. Population structure and energetics of the bopyrid isopod parasite Orthione Griffenis in mud shrimp Upogebia Pugettensis. Journal of Crustacean Biology. 2008 Apr 1;28(2):228–33.

24. Abel RL, Laurini CR, Richter M.). A palaeobiologist’s guide to ’virtual’ micro-CT preparation. Palaeontologica Electronica. 2012 Apr 29;15(2):1–17.

25. Hounsfield GN. Computerized transverse axial scanning (tomography). 1. Description of system. Br. J. Radiol. 1973 Dec;46(552):1025–88.

26. McCormack AM. Representation of a function by its line integrals, with some radiological applications. J. Appl. Phys. 1963 Sep;34(9):2722–27.

27. Conroy GC, Vannier MW. Noninvasive three-dimensional computer imaging of matrix-filled fossil skulls by high-resolution computed tomography. Science. 1984 Oct 26;226(4673):456–58.

28. Sutton M, Rahman I, Garwood R. Virtual paleontology – an overview. The Paleontological Society Papers. 2017 Apr 27;22:1–20.

29. Klompmaker AA, Kloess PA, Jauvion C, Brezina J, Landman NH. Internal anatomy of a brachyuran crab from a Late Cretaceous methane seep and an overview of internal soft tissues in fossil decapod crustaceans. Palaeontologia Electronica. 2023 Sep 23;26.3:a44.

30. Luque J, Xing L, Briggs DEG, Clark EG, Duque A, Hui J, Mai H, McKellar RC. Crab in amber reveals an early colonization of nonmarine environments during the Cretaceous. Science Advances. 2021 Oct 20;7(43): eabj5689.

31. Waters JA, White LE, Sumrall CD, Nguyen BK. A new model of respiration in blastoid (Echinodermata) hydrospires based on computational fluid dynamic simulations of virtual 3D models. Journal of Paleontology. 2017 May 10;91(4):662–71.

32. Bicknell RDC, Ledogar JA, Wroe S, Gutzler BC, Watson III WH, Paterson JR. Computational biomechanical analyses demonstrate similar shell-crushing abilities in modern and ancient arthropods. Proc. R. Soc. B. 2018 Oct 24;285:20181935.

33. Zhang Y, Xu D, Li J, Zhang Z, Ding S, Wu W, Xia R. Mechanical properties and clamping behaviors of snow crab claw. Journal of the Mechanical Behavior of Biomedical Materials. 2021 Dec;124:104818.

34. Bright JA. A review of paleontological finite element models and their validity. Journal of Paleontology. Cambridge University Press. Jul 2014;88(4):760–9.

35. Rayfield EJ. Finite element analysis and understanding the biomechanics and evolution of living and fossil organisms. Annur. Rev. of Earth and Planetary Sciences. 2007 May 30;35:541–76.

36. Cunningham JA, Rahman IA, Lautenschlager S, Rayfield EJ, Donoghue PCJ. A virtual world of paleontology. Trends Ecol. Evol. 2014 Jun;29(6):347–57.

37. Smith SI. Preliminary report on the Brachyura and Anomura dredged in deep water off the south coast of New England by the United States Fish Commission in 1880, 1881, and 1882. Proceedings of the United States National Museum. 1883 Jun 18;6(1):1–57.

38. Collins JSH, Rasmussen HW. Upper Cretaceous – Lower Tertiary decapod crustaceans from West Greenland. Bulletin Grønlands Geologiske Undersøgelse. 1992 Jan 1;162:1–46.

39. Portell RW, Collins JSH. Decapod crustaceans of the Lower Miocene Montpellier Formation, White Limestone Group of Jamaica. In. Donovan, S.K. (ed.) The Mid-Cainozoic White Limestone Group of Jamaica. Cainozoic Research. 2004;31(1-2):109–26.

40. North Star Imaging. SubpiX: An advanced CT scanning technique. 2021 Oct 9. https://4nsi.com/subpix-an-advanced-ct-scanning-technique/

41. xComet Technologies Canada Inc. Object Research Systems (ORS) Inc. Dragonfly (Version 2022.1). Updated 2022 Feb 14. https://www.theobjects.com/dragonfly/index.html.

42. Blender Foundation. Blender - a 3D modelling and rendering package (Version 3.4). 2003. Updated 2022 Dec 7. http://www.blender.org

43. Zienkiewicz OC, Taylor RL, Zhu JZ. The Finite Element Method: Its Basis and Fundamentals. Elsevier Butterworth-Heinemann. 2005.

44. Riegel J, Mayer W, van Havre Y. FreeCAD (version 0.20.2). 2002 Oct 29. Updated 2022 Dec 6. https://www.freecad.org/

45. Geuzaine C, Remacle JF. Gmsh: A 3-D finite element mesh generator with built-in pre- and post-processing facilities. International journal for numerical methods in engineering. 2009 Sep 10;79(11):1309–31.

46. Chen P, Lin AY, McKittrick J, Meyers MA. Structure and mechanical properties of crab exoskeletons. Acta Biomateriala. 2008 May;4(3):587–96.

47. Gadgey KK, Bahekar A. Investigation of mechanical properties of crab shell: a review. International Journal of Latest Trends in Engineering and Technology. 2017;8(1):268–81.

48. Amato CG, Waugh DA, Feldmann RM, Schweitzer CE. Effect of calcification on cuticle density in decapods: A key to lifestyle. Journal of Crustacean Biology. 2008 Oct 1;28(4):587–95.

49. Kim SS, Kim SJ, Moon YD, Lee YM. Thermal characteristics of chitin and hydroxypropyl chitin. Polymer. 1994 Jul;35(15):3212–16.

50. Sachs C, Fabritius H, Raabe D. Experimental investigation of the elastic–plastic deformation of mineralized lobster cuticle by digital image correlation. Journal of Structural Biology. 20006 Jun 21;155:409–25.

51. Swanson BO, George MN, Anderson SP, Christy JH. Evolutionary variation in the mechanics of fiddler crab claws. BMC Evol Biol. 2013 Jul 15;13:137.

52. Blundon JA, Kennedy VS. Mechanical and behavioral aspects of blue crab, Callinectes sapidus (Rathbun), predation on Chesapeake Bay bivalves. Journal of Experimental Marine Biology and Ecology. 1982 Nov 5;65(1):47–65.

53. Govind CK, Blundon JA. Form and function of the asymmetric chelae in blue crabs with normal and reversed handedness. Biological Bulletin. 1985 Apr;168(2):321–31.

54. Dhondt G, Wittig K. CALCULIX: A free software three-dimensional structural finite element program. 1998. Updated 2016 May 8. http://www.calculix.de/

55. Kitware, Los Alamos National Laboratory. Paraview (version 5.11.0). 2002. Updated 2022 Nov 17. https://www.paraview.org/

56. Taylor GI, Quinney H. The plastic distortion of metals. Phil Trans R Soc A. 1931 Nov 13;230:323–62.

57. National Evolutionary Synthesis Center. Dryad Digital Repository. 2008 Jan. https://datadryad.org/stash

58. Carlson CJ, Burgio KR, Dougherty ER, Phillips AJ, Bueno VM, Clements CF, Castaldo G, Dallas TA, Cizaukas CA, Cumming GS, Doña J, Harris NC, Jovani R, Mironov S, Muellerklein OC, Proctor HC, Getz WM. Parasite biodiversity faces extinction and redistribution in a changing climate. Science Advances. 2017 Sep 6;3(9):e1602422.

59. Cizauskas CA, Carlson CJ, Burgio KR, Clements CF, Dougherty ER, Harris NC, Phillips AJ. Parasite vulnerability to climate change: an evidence-based functional trait approach. R. Soc. Open sci. 2017 Jan 1;4:160535.

60. Byers JE. Effects of climate change on parasites and disease in estuarine and nearshore environments. PLoS Biol. 2020 Nov 24;18(11):e3000743.

61. Wood CL, Welicky RL, Preisser WC, Leslie KL, Mastick N, Greene C, Maslenikov KP, Tornabene L, Kinsella JM, Essington TE. A reconstruction of parasite burden reveals one century of climate-associated parasite decline. PNAS. 2023 Jan 9;120(3):e2211903120.

62. Short EE, Caminade C, Thomas BN. Climate change contribution to the emergence or re-emergence of parasitic diseases. Infect Dis. 2017 Sep 25;10:1178633617732296.

63. McInerney FA, Wing SL. The Paleocene-Eocene thermal maximum: A perturbation of carbon cycle, climate, and biosphere, with implications for the future. Annu. Rev. Earth Planet. Sci. 2011 Mar 1;39:489–516.

64. Luquet G. Biomineralizations: insights and prospects from crustaceans. Zookeys. 2012 Mar 20;176:103–21.

65. Marcé-Nogué J. One step further in biomechanical models in palaeontology: a nonlinear finite element analysis review. PeerJ. 2022 Aug 8;10:e13890.

